## Supplementary Figures for "Establishing the green algae *Chlamydomonas incerta* as a platform for recombinant protein production"

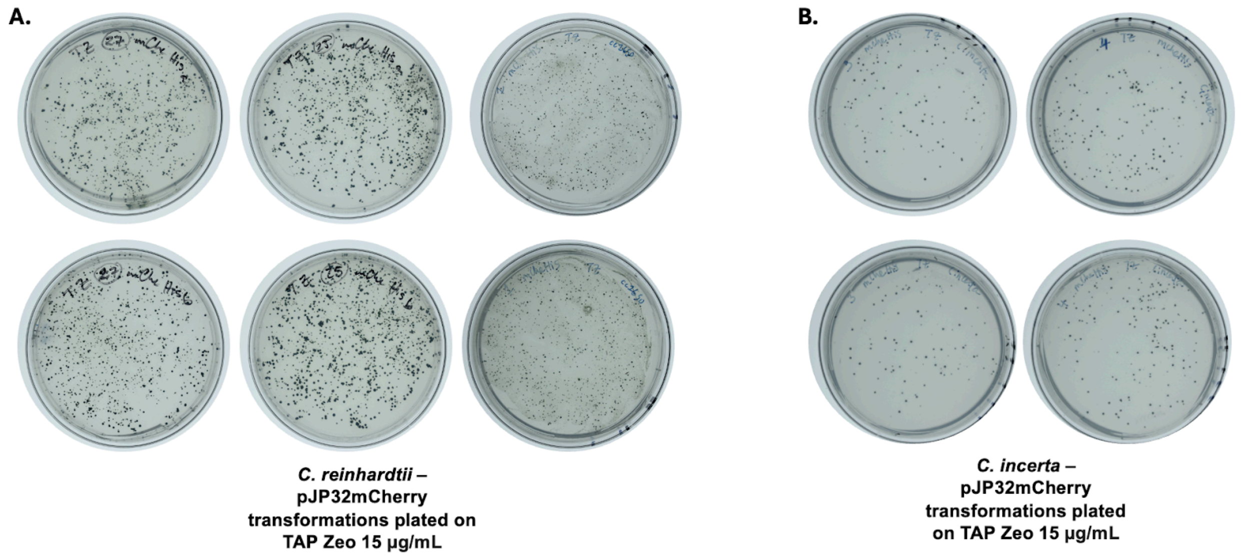


**Supplementary Figure 1. Transformations of pJP32mCherry into *C. reinhardtii* and *C. incerta*.**

The mCherry secretion vector, pJP32mCherry, was transformed into **A)** *C. reinhardtii* and **B)** *C. incerta*. Triplicate transformations were performed for both species, and transformants were spread onto two plates (paired vertically). However, only two transformations for *C. incerta* generated an adequate amount of colonies, so the third transformation is not shown.

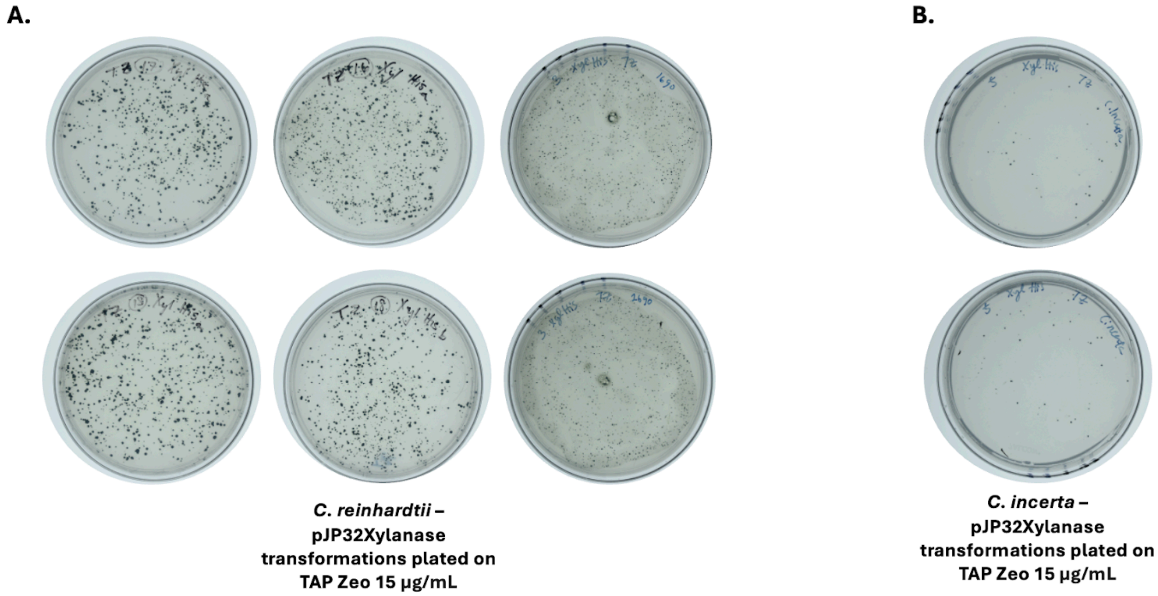


**Supplementary Figure 2. Transformations of pJP32Xylanase into *C. reinhardtii* and *C. incerta*.**

The xylanase secretion vector, pJP32Xylanase, was transformed into **A)** *C. reinhardtii* and **B)** *C. incerta*. Triplicate transformations were performed for both species, and transformants were spread onto two plates (paired vertically). However, only one transformation for *C. incerta* generated an adequate amount of colonies, so the other two transformations are not shown.

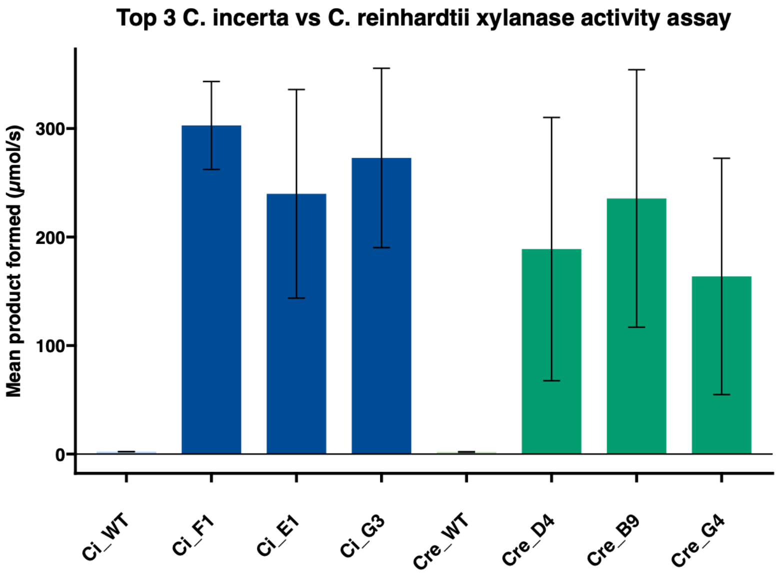


**Supplementary Figure 3. Xylanase activity assay of three highest expressing *C. incerta* and *C. reinhardtii* transgenic lines of pJP32Xylanase.**

Hydrolysis of the fluorogenic substrate 6,8-difluoro-4-methylumbelliferyl β-d-xylobioside (DiFMUX2) by xylanase led to increased fluorescence at an excitation wavelength of 385 nm and emission wavelength of 455 nm over time. The F1, E1, and G3 strains of transgenic *C. incerta* expressing xylanase formed 302.799 µmol/s, 239.806 µmol/s, and 272.852 µmol/s of product, respectively. The D4, B9, and G3 strains of transgenic *C. reinhardtii* expressing xylanase formed 188.873 µmol/s, 235.523 µmol/s, and 163.695 µmol/s of product, respectively. The *C. incerta* and *C. reinhardtii* wild types formed 2.219 µmol/s and 1.918 µmol/s of product, respectively.

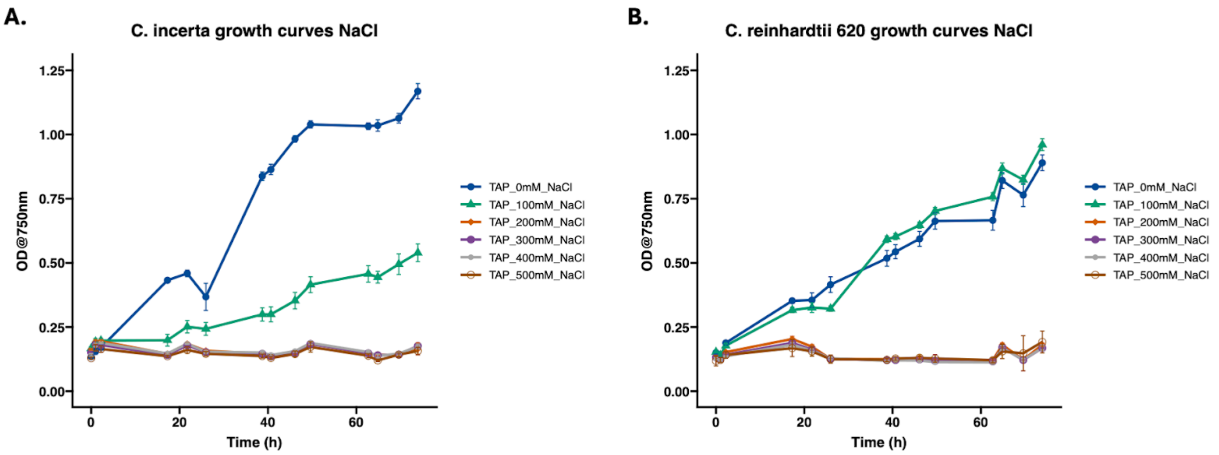


**Supplementary Figure 4. Growth curves of *C. incerta* and *C. reinhardtii* in TAP media with NaCl.**

**A)** The *C. incerta* wild type was grown in TAP media containing 0 mM, 100 mM, 200 mM, 300 mM, 400 mM, and 500 mM NaCl. Absorbance readings at 750 nm were measured using the Infinite® M200 PRO plate reader (Tecan, Männedorf, Switzerland) over approximately 73 hours, and the readings represent the average of biological quadruplicates. **B)** The *C. reinhardtii* wild type was grown in TAP media containing 0 mM, 100 mM, 200 mM, 300 mM, 400 mM, and 500 mM NaCl. Absorbance readings at 750 nm were measured using the Infinite® M200 PRO plate reader (Tecan, Männedorf, Switzerland) over approximately 73 hours, and the readings represent the average of biological quadruplicates.

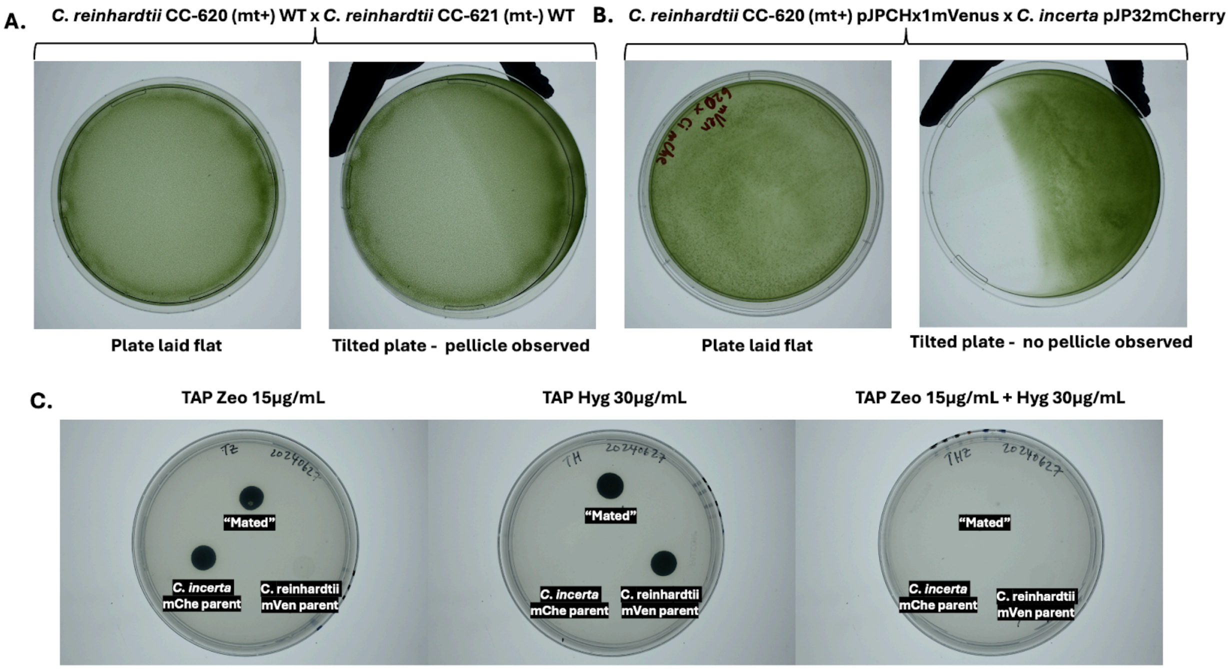


**Supplementary Figure 5. Unsuccessful interspecies mating between *C. incerta* and *C. reinhardtii*.**

**A)** *C. reinhardtii* CC-620 (mt+) wild type and *C. reinhardtii* CC-621 (mt-) wild type were mated, and a pellicle phenotype was observed after 12-16 hours, indicating success of intraspecies mating. **B)** Transgenic *C. reinhardtii* CC-620 (mt+) pJPCHx1mVenus and transgenic *C. incerta* pJP32mCherry, and a pellicle phenotype was not observed after 12-16 hours, indicating that interspecies mating is not feasible. **C)** The mixed cells that did not formed a pellicle from B, along with the *C. reinhardtii* pJPCHx1mVenus and *C. incerta* pJP32 parents were plated onto 3 types of plates: TAP agar plates containing zeocin 15 µg/mL, TAP agar plates containing hygromycin B 30 µg/mL, and TAP agar plates containing both zeocin 15 µg/mL and hygromycin 30 µg/mL.

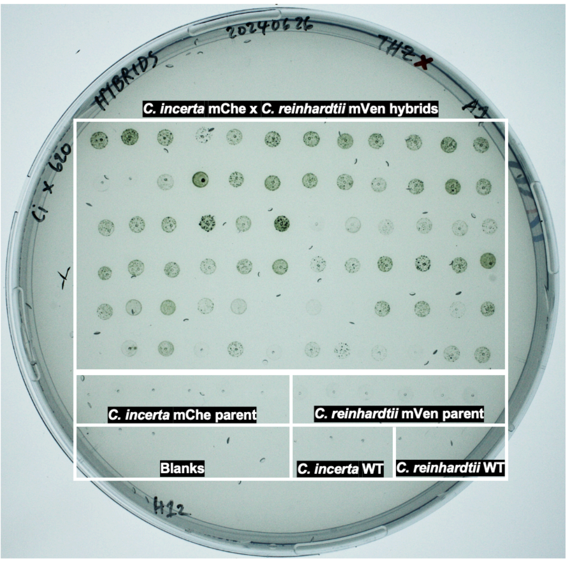


**Supplementary Figure 6. Screening for successful hybridizations between transgenic *C. incerta* pJP32mCherry and transgenic *C. reinhardtii* CC-620**

Potential hybrid colonies from the transformation plate were picked into 96-well plates containing TAP media and grown for 7 days. Replica TAP agar plate containing zeocin 15 μg/mL and hygromycin 30 μg/mL was made to screen for stable hybrids.

| Gene | GenBank ID | Best hit GenBank Protein ID | E-value | Query coverage (%) | Identity (%) |
|---|---|---|---|---|---|
| rbcs | X04472.1 | KAG2425659.1 | 1e-130 | 100 | 95.14 |
| hsp70 | M76725.2 | KAG2445402.1 | 0.0 | 100 | 98.31 |
| maw8 | XM_043064089 (NCBI reference sequence) | KAG2440857.1 | 1e-168 | 84 | 86.26 |
| gp1 | AF309494.1 | ABK42021.1 | 5e-105 | 31 | 80.77 |

**Supplementary Table 1. BLASTp results comparing protein sequences of genes with DNA parts in the  between *C. incerta* and reference protein sequences.**

The table shows the GenBank IDs for the queried genes, the best hit GenBank protein ID, the E-value indicating the statistical significance of the match, the query coverage percentage, and the percentage identity of the aligned sequences.

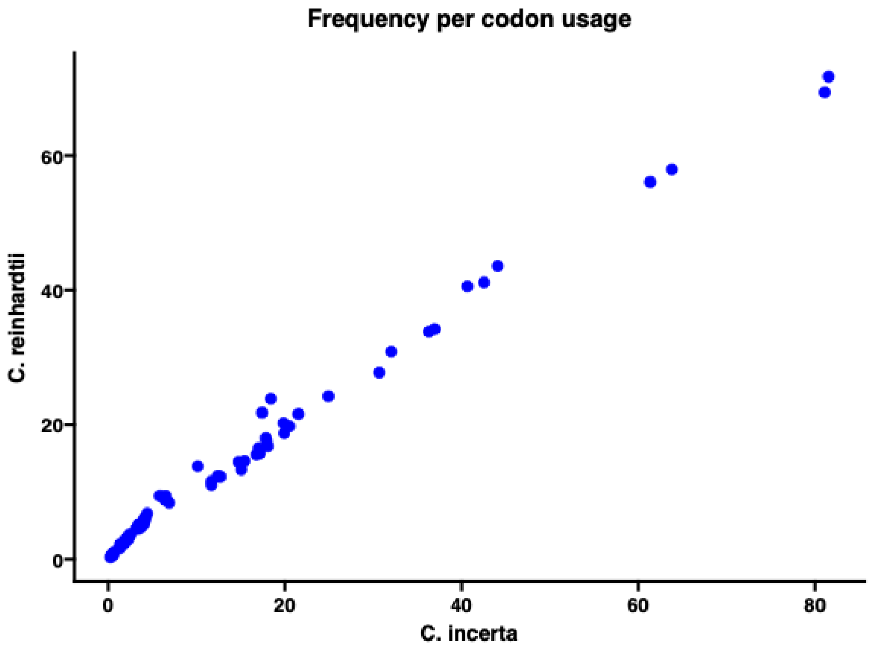


**Supplementary Figure 7. Codon usage comparison between *C. incerta* and *C. reinhardtii.***

The scatter plot depicts the frequency of each codon's usage in *C. incerta* (x-axis) against *C. reinhardtii* (y-axis). A positive 1:1 correlation in codon usage patterns is observed between the two species, suggesting similarities in their codon preferences despite genomic differences.

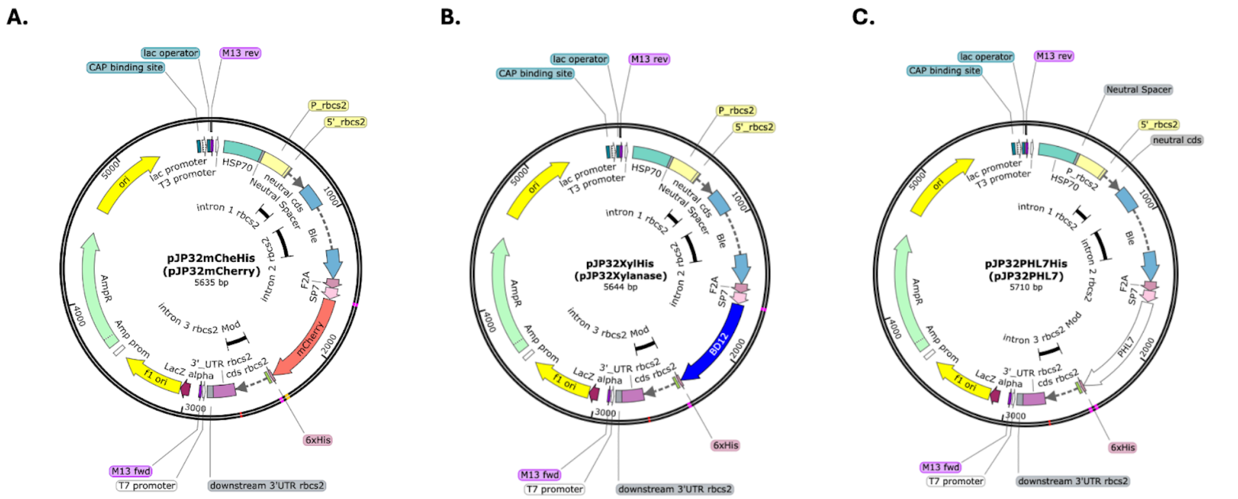


**Supplementary Figure 8. Plasmid maps for pJP32 secretion vectors.**

Plasmid maps for **A)** pJP32mCheHis (denoted as pJP32mCherry in the publication), **B)** pJP32XylHis (pJP32Xylanase), and **C)** pJP32PHL7His (pJP32PHL7) using SnapGene (GSL Biotech LLC, San Diego, CA, USA).

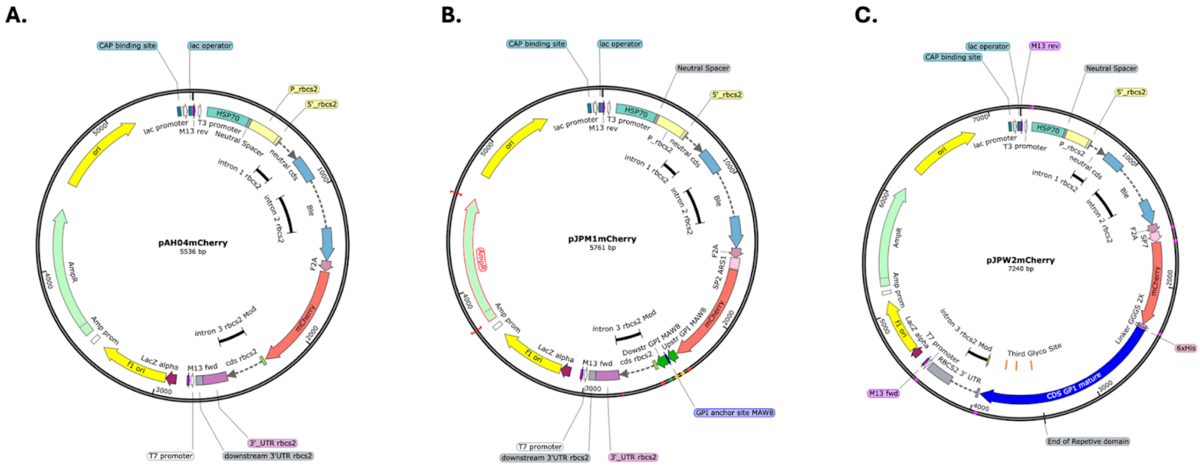


**Supplementary Figure 9. Plasmid maps for pAH04, pJPM1, and pJPW2 vectors.**

Plasmid maps for **A)** pAH04mCherry, an mCherry cytosolic expression vector, **B)** pJPM1mCherry, an mCherry cell membrane expression vector, and **C)** pJPW2mCherry, an mCherry cell wall expression vector, using SnapGene (GSL Biotech LLC, San Diego, CA, USA).

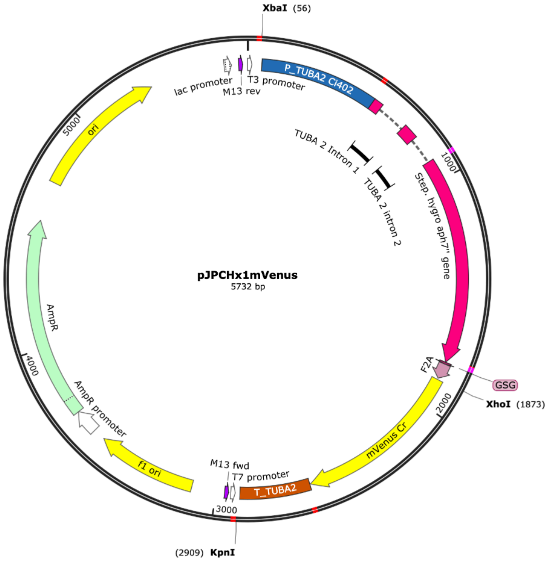


**Supplementary Figure 10. Plasmid map for pJPCHx1mVenus vector.**

Plasmid map for pJPCHx1mVenus, an mVenus cytosolic expression vector containing the hygromycin B antibiotic resistance gene, using SnapGene (GSL Biotech LLC, San Diego, CA, USA).

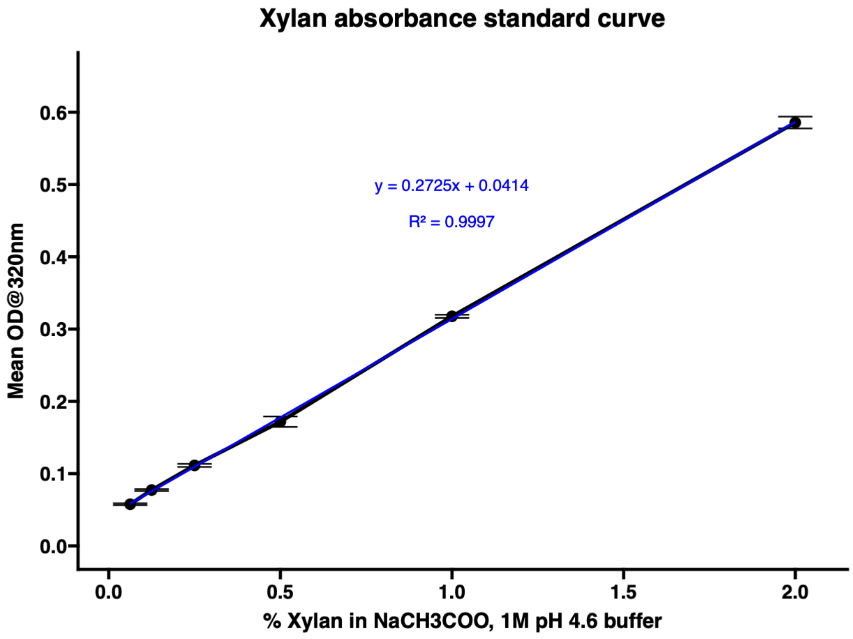


**Supplementary Figure 11. Standard curve of absorbance of Xylan from Corn Core.**

The xylan absorbance standard curve was created using varying concentrations of Xylan from Corn Core (0%, 0.0625%, 0.125%, 0.25%, 0.5%, 1%, and 2% w/v) in sodium acetate buffer (1M, pH 4.5). Absorbance at 320 nm were measured in the 96-well UV-Star® microplates (Greiner Bio-One, Kremsmünster, Austria) using the Infinite® M200 PRO plate reader (Tecan, Männedorf, Switzerland). The standard curve was fitted with a linear regression equation y = mx + b, where m is the slope and b is the y-intercept (equation: y = 0.2727x + 0.0414; $R^2$ = 0.9997).
